# Structural landscape of base pairs containing post-transcriptional modifications in RNA

**DOI:** 10.1101/098871

**Authors:** Preethi S. P., Purshotam Sharma, Abhijit Mitra

## Abstract

Base pairs involving post-transcriptionally modified nucleobases are believed to play important roles in a wide variety of functional RNAs. Here we present our attempts towards understanding the structural and functional role of naturally occurring modified base pairs by analyzing their distribution in different RNA classes, with the help of crystal structure and sequence database analyses. In addition, we quantify the variation in geometrical features of modified base pairs within RNA structures, and characterize their optimum geometries and binding energies using advanced quantum chemical methods. Further comparison of modified base pairs with their unmodified counterparts illustrates the effect of steric and electronic structure alterations due to base modifications. Analysis of specific structural contexts of modified base pairs in RNA crystal structures revealed several interesting scenarios, including those at the tRNA:rRNA interface, antibiotic-binding site and the three-way junctions within tRNA, which when analyzed in context of available experimental data, allowed us to correlate the occurrence and strength of modified base pairs with the specific functional roles they play in context of RNA macromolecules.

## INTRODUCTION

Recent structural and mechanistic studies on RNA molecules illustrate that although tremendous progress has been achieved towards understanding their versatile role in different aspects of modern biology, there is a need to deepen our understanding of the principles governing the structure, dynamics and functions of these fascinating biomacromolecules. For example, similar to proteins, where the post-translational modifications are associated with catalysis, initiation and termination of signal cascades, and integration of information at many metabolic intersections (Walsh et al. 2005), post-transcriptionally modified nucleobases may also be associated with a variety of RNA functionalities. A detailed understanding of chemical modifications of RNA nucleobases, and resulting changes in associated noncovalent interactions, is therefore one of the necessary requirements for investigating the functional diversity of RNA molecules.

Posttranscriptional modifications in RNA range from the addition of simple functional groups (e.g. base/ribose methylation) to complex side chains (e.g. hypermodifications) (Denmon et al. 2011). In addition, such modifications may also include substitutions (e.g. conversion of uridine to 4-thiouridine, s^4^U), isomerization (e.g. conversion of uridine to pseudouridine, Ψ) and reduction (e.g. conversion of uridine to dihydrouridine D, Fig. 1, (Mueller et al. 1998)). Survey of available literature suggest that nucleobase modifications are known to serve as important evolutionary tool for tuning up the RNA structure to perform its biological functions with greater fidelity (Emmerechts et al. 2008). Thus, it is not surprising that chemically modified ribonucleotides are present in RNA of organisms belonging to all three (i.e. archaea, bacteria and eukarya) domains of life (Decatur and Fournier 2002), where the percentage of chemical modifications in a RNA sequence is roughly proportional to the complexity of the organism (Chow et al., 2007). In this context, repositories of modified RNA bases available in databases such as MODOMICS (Dunin-Horkawicz et al. 2006) and RNAMDB (Cantara et al. 2011) provide a comprehensive listing of post-transcriptionally modified nucleosides in RNA, which are useful in understanding RNA nucleoside modification pathways.

**FIGURE 1.**
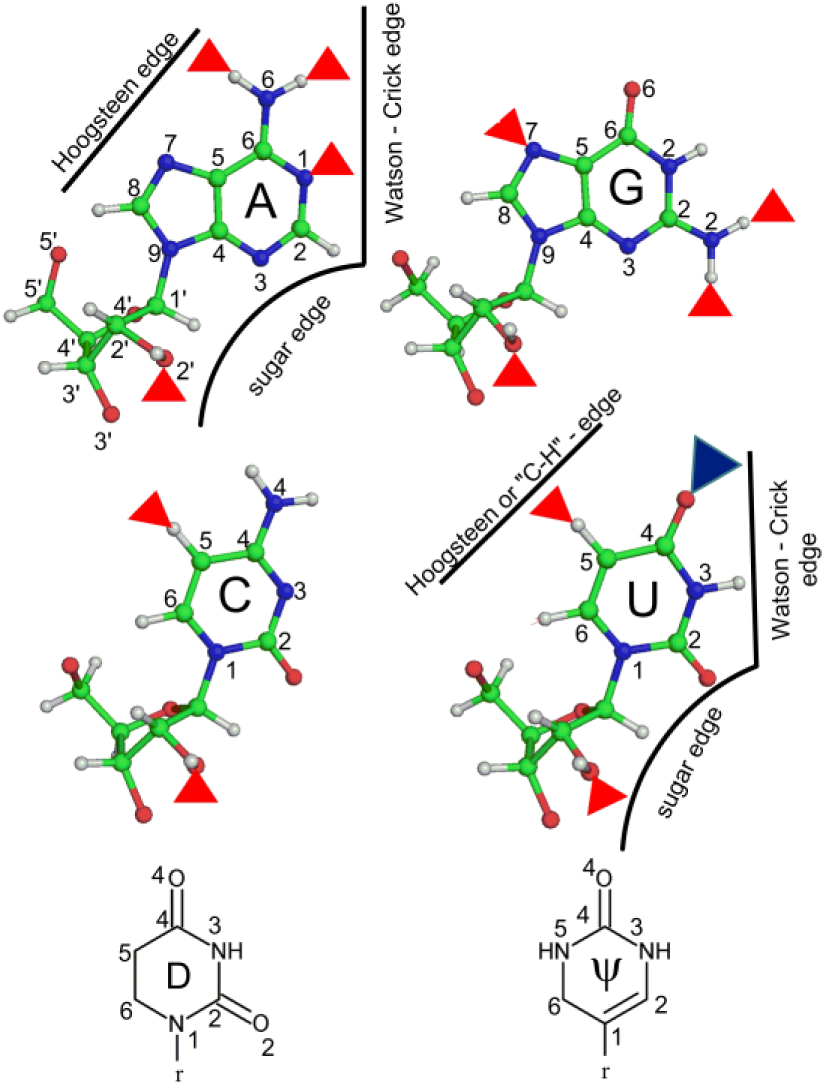
Schematic representation of modified base pairs showing their interacting edges. Red triangles represent modification involving methyl group substitution, whereas blue triangle represents substitution of oxygen with sulfur atom. The ribose sugar is represented by r in the structures of dihydrouridine (D) and pseudouridine (ψ).

**FIGURE 2.**
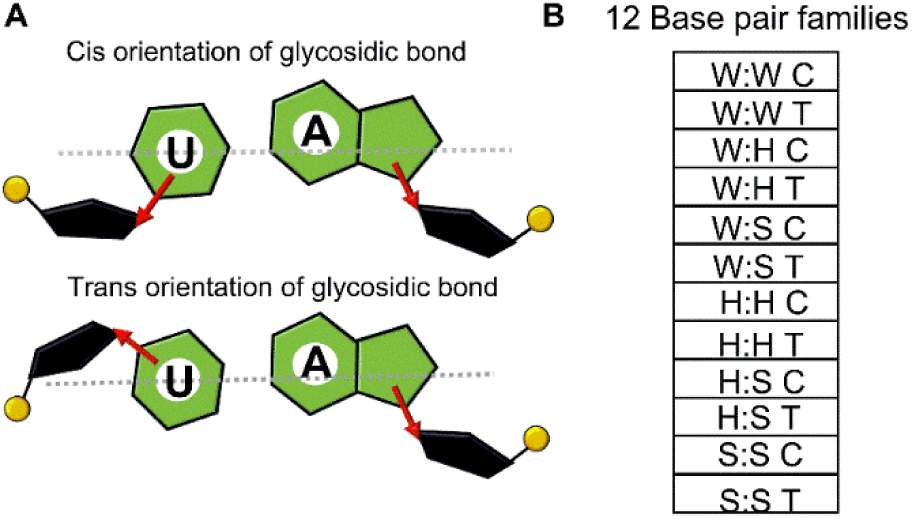
(A) Schematic representation of cis (C) or trans (T) orientation of the glycosidic bond. (B) List of 12 RNA base pairing families. W, H and S represent Watson-Crick, Hoogsteen and sugar edges respectively.

In terms of mechanistic understanding, one of the ways through which modified nucleobases may provide stability to the RNA tertiary structures is by inducing tailor-made alterations to the conformational preferences of corresponding nucleotides. For example, methylation at the 2’–OH group of ribose shifts the equilibrium towards C3’-endo sugar pucker, thus favoring the A-form RNA helices (Motorin and Helm 2010). Further, contrary to the naturally occurring nucleotides which adopt the *anti* conformation, pseudouridine prefers the *syn* conformation at the glycosidic bond. Given the low energy requirement for the *anti/syn* transition, pseudouridine can shift between the two conformations with relatively greater ease, and can function as a conformational switch in RNA (Charette and Gray 2000). In addition, dihydrouridine significantly destabilizes the C3’-*endo* sugar conformation, which is associated with base stacked, ordered, A-type helical RNA (Dalluge et al. 1996). Thus, it is not surprising that dihydrouridine is found in higher percentage in organisms that grow in low temperatures (psychrophiles), where it allows for extra flexibility for RNAs that function near the freezing point of water (Dalluge et al. 1996).

Apart from changing the nucleotide conformational preferences, chemical modifications can also affect the noncovalent interactions involving nucleobases in RNA (Davis, 1995). Base pairing and base stacking constitute the major noncovalent interactions through which nucleotides interact with each other in RNA. Although base stacking provides the driving force for RNA folding, it is relatively weaker and less specific compared to base pairing (Leontis and Westhof 2001). On the other hand, base pairing being a relatively stronger force, provides directionality and specificity (Leontis and Westhof 2001; Leontis et al. 2002), and plays a crucial role in scripting the structural variety and functional dynamics of RNA molecules. Although tRNA base pairing classification efforts by Leontis, Westhof and others (Stombaugh et al. 2009), quantum chemical revelation of physicochemical principles of RNA base pairing (Šponer et al. 2005a; Šponer et al. 2005b; Šponer et al. 2005c; Sharma et al. 2008; Sharma et al. 2010a; Sponer et al. 2010; Chawla et al. 2011; Halder et al. 2014; Halder et al. 2015) and increasing availability of X-ray crystal structures of functional RNA molecules have significantly enhanced our understanding of base pairing interactions involving canonical nucleosides in RNA, the effect of chemical modifications of nucleosides on intrinsic changes in RNA base pairs been considered only in a few quantum chemical or structural studies (Oliva et al. 2006; Oliva et al. 2007; Chawla et al. 2015).

Previous studies on tRNA post transcriptional modifications observed that modifications that introduce positive charge strongly stabilize the geometry of the corresponding base pairs. An example is the stabilization of a reverse Watson-Crick geometry of G15:C48 tertiary interaction in RNA on positively charged archaeosine modification of guanine (Oliva et al. 2007). More recently, analysis of available RNA crystal structures (Chawla et al. 2015) revealed that 11 types of base modifications participate in base pair formation, forming 27 distinct base pair combinations. Further quantum chemical studies revealed that whereas methyl modifications either impart steric clashes or introduce positive charge, other modifications such as Ψ and D affect the stability and flexibility of the structure (Chawla et al. 2015).

Be that as it may, there remains a significant gap in understanding of the structural principles involving chemically modified base pairs in RNA, and, a number of factors need to be considered, in order to address it. First, since the structural diversity of modifications varies across different groups of RNAs (Cantara et al. 2011), the relative abundance of modified base pairs with respect to different RNA classes need to be considered. Further, due to the prevalence of sugar modifications in RNA, and given the fact that ribose sugar plays an important role in RNA base pairing (Šponer et al. 2005d; Sharma et al. 2008; Mládek et al. 2009), the effect of sugar modifications on geometries and stabilities of RNA base pairs need to be analyzed. In addition, the geometrical characteristics of crystal occurrences of modified base pairs need to be analyzed in detail, in order to quantify the effect of base modifications on conformational flexibilities of base pairs in crystal contexts. Furthermore, the structural context of occurrence of modified base pairs in RNA structures need to be analyzed in detail, in order to evolve deeper understanding of the functional roles of such base pairs.

In the present work, we attempt to fill this void in literature by probing into the geometrical features and the intrinsic stabilities of base pairs containing modified RNA bases, in terms of their molecular-level interactions, as well as their macromolecular context of occurrence in RNA structures. For this, we have chosen a multipronged approach that employs a combination of sequence analysis, crystal structure database analysis using tools of structural bioinformatics, as well as state-of-the-art quantum chemical methods. Overall, our study provides a comprehensive analysis of modified base pairs in RNA, which may inspire future studies on the specific functional context of individual base modifications in RNA.

## RESULTS AND DISCUSSION

### Sequence – structure – energetic context of occurrences of modified base pairs in functional RNAs

#### Statistical overview of modified base pairs in RNA 3D structures

i. *tRNA structures show a remarkably high occurrence of modified base pairs*: Fifteen different types of naturally occurring modified RNA nucleosides were searched to analyze their propensities to form base pairs (Table 1). 11 of them involved modification(s) of the nucleobase moiety, and were previously found to participate in base pairing in RNA structures (Chawla et al. 2015). Since methylation of 2’–OH group is also known to affect base pairing through alteration of sugar edge interactions of RNA bases (Leontis and Westhof 2001), base pairs involving methylation at the 2’–OH group of ribose sugar were also considered for all four nucleosides (A, C, G and U). A set of 207 high-resolution RNA crystal structures containing at least one modified base (Supplemental Table S1), was selected for analysis according to specific search criteria (see Materials and Methods). 80% of the crystal structures belong to four major RNA classes (tRNA (25%), 16S rRNA (24%), 23S rRNA (23%) and RNA binding proteins (11%), Fig. 3A). However, the occurrence of modified bases in rest of RNA classes was rather marginal (each below 10%). 65% (135) of the total (207) crystal structures contained at least one modified base that participates in base pairing (Supplemental Table S2). Approximately one third of such structures belong to tRNA (36%), another one third belong to 23S rRNA (34%) and one-sixth belong to 16S rRNA (18%, Fig. 3B). More than half of the modified bases within the dataset participate in base pairing, whereas the unpaired modified bases were present in other variable structural contexts (Supplemental Tables S2-S4). A total of 453 modified base pairs were detected from these RNA crystal structures. half of which belong to tRNA (Supplemental Fig. S1A). This is in synchrony with previous studies that observed relatively greater occurrence of modified bases in tRNA compared to other RNA classes (Limbach et al. 1994; Machnicka et al. 2014). Given the fact that most of the nucleobases in tRNA are involved either in base pairing or in tertiary interactions (Oliva et al. 2006), it is not surprising that most of the modified bases present in tRNA also participate in base pairing.
ii. *Base pairs containing modified uridine or guanosine are relatively more abundant*: Crystal structure analysis reveals that 72% of the modified base pairs contained either uridine (37%) or guanosine (35%) modification (Supplemental Fig. S1B). The greater proportion of base pairs containing uridine modifications can be attributed to the natural occurrence of a rich variety in uridine modifications (base methylation and/or sugar methylation, thiolation, pseudouridylation or reduction), each of which has the propensity to form base pairs. In fact, 6 of the 15 modified nucleosides that form base pairs (Table 1), contain modification of uridine. On the other hand, greater abundance of base pairs containing guanosine modifications can be correlated to occurrence of a variety of methylation sites at guanosines (e.g. N1, N2, N7 or 2’–OH), as well as the propensity of guanosine to form singly and doubly methylated structures at N2, all of which participate in base pairing.
iii. *Methylation is the preferred chemical modification in RNA base pairs*: Distribution of modified base pairs with respect to the type of modification reveals that more than half of them contain at least one methylated base (60% total, 35% in tRNA and 22% in 16S rRNA, Further, substantial diversity is observed in methylated base pairs, where the relative population depends on the parent nucleobase. For example, one third of the total methylated base pairs contain m^5^C, followed by methylated G (26%) that include m^7^G (13%), m^2^G (9%) and m^2^_2_G (4%, Supplemental Fig. S1C). However, in contrast to the abundance (86%) of base pairs containing base modifications, only 14% of modified base pairs contain sugar methylation at 2’-OH (Supplemental Fig. S1C). The greater abundance of methylated base pairs in RNA structures can be correlated to the wide variety of structural roles played by these bases that include enhancement of base stacking and increase in nucleobase polarizability by base methylations, and tendency to favor C3’ endo–conformation, block sugar-edge interactions and enhancement of stability against hydrolysis by methylation at 2’-OH of sugar (Helm 2006).
iv. *Modified base pairs are observed in all major RNA structural elements and span diverse RNA base pairing geometries*: Distribution of modified base pairs with respect to their contextual occurrence in RNA crystal structures reveals that approximately half (49%) of them are present in stem (helical) regions, 14% in loop regions and rest 37% are involved in tertiary interactions Supplemental Fig. S1D). Overall, the results are in line with a previous crystal structure analysis (Chawla et al. 2015), where 41% of the total modified base pairs were found to be involved in tertiary interactions. Categorization of modified base pairs in terms of the portion of the nucleoside that interacts with the partner nucleoside reveals that 80% of them involve base-base (B-B) interactions, 12% involve base-nucleoside (B-S) interactions and 8% involve nucleoside-nucleoside (S-S) interactions (Supplemental Table S7).Further categorization of base pairs in terms of the interacting edge (Watson-Crick (W), Hoogsteen (H) or Sugar (S)) and the glycosidic bond (cis or trans, Figure 1) orientation reveals that the B-B interactions involving modified bases (80%) span four of the six associated base pairing families – W:WC (49%), W:HT (27%), W:WT (3%) and W:HC (1%). Notably, no examples of modified base pairs are observed among H:HC and H:HT families of base pairs, mainly because, owing to their unique backbone topology requirements, base pairs involving H:H occur rarely in RNA structures (Sharma et al. 2010a). Thus, the possibility of base modifications in H:H base pairs is rare. On the other hand, B-S (12%) and S:S (8%) interactions span all six possible base pair geometries (W:SC (5%), W:ST (4%), H:ST (2%), H:SC (1%) S:SC (6%) and S:ST (2%, Supplemental Tables S7). Overall, the total 453 base pairs identified in RNA crystal structures belong to 36 unique base pairing combinations, 24 of which involve B-B interactions, 6 involve B-S interactions and 6 involve S:S interactions (Table 2).

**FIGURE 3.**
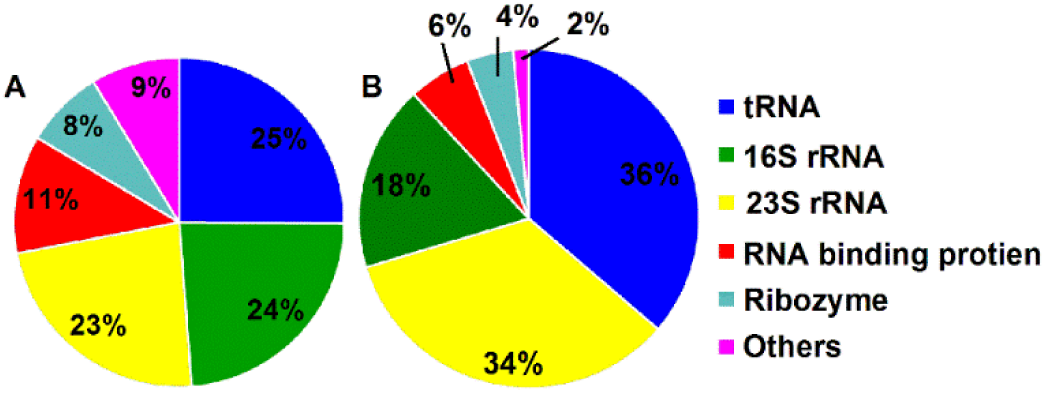
(A) Percent distribution of total 207 crystal structures in the dataset as a function of RNA type. (B) Percent distribution of those 135 crystal structures as a function of RNA type that contain at least one modified base.

**TABLE 1.** Naturally occurring modified bases that participate in base pairing in RNA crystal structures.

| Modified bases<sup>a</sup> | Observed base pairs |
| --- | --- |
| N1-methyl adenine (m<sup>1</sup>A) | m<sup>1</sup>A:A W:HT, m<sup>1</sup>A:m<sup>5</sup>U<br>W:HT, m<sup>1</sup>A:m<sup>5</sup>Um W:HT |
| N6,6-dimethyl adenine (m<sup>6</sup><sub>2</sub>A) | m<sup>6</sup><sub>2</sub>A:G S:ST |
| 2' -O-methyl adenine (Am) | Am:G S:ST |
| N2-methyl guanine (m<sup>2</sup>G) | m<sup>2</sup>G:C W:WC, m<sup>2</sup>G: C<br>W:W(+ )C, m<sup>2</sup>G:U W:WC |
| N2,2-dimethyl guanine (m<sup>2</sup><sub>2</sub>G) | m<sup>2</sup><sub>2</sub>G:A W:WC |
| N7-methyl guanine (m<sup>7</sup>G) | m<sup>7</sup>G:G W:HT, m<sup>7</sup>G:C<br>W:WC, m<sup>7</sup>G:A S:ST,<br>m<sup>7</sup>G:A S:WT |
| 2'-O-methyl guanine (Gm) | Gm:A W:WC, Gm:G<br>S:SC, Gm:C W:WC, |
| C5-methyl cytosine (m<sup>5</sup>C) | m<sup>5</sup>C:G W:WC, m<sup>5</sup>C:G<br>W:WT |
| 2'-O-methyl cytosine (Cm) | Cm:Gm W:WC |
| Pseudouridine (Ψ) | Ψ:A W:WC, Ψ:A W:HT,<br>Ψ:G W:WC, Ψ:C S:WC |
| 5,6-dihidrouridine (D) | D:G W:ST, D:G H:SC,<br>D:U W:WT, D:U S:ST |
| C5-methyl uracil (m<sup>5</sup>U) | m<sup>5</sup>U:A W:HT, m<sup>5</sup>U:G<br>W:WC, m<sup>5</sup>U:G W:HT |
| 2',5-dimethyl uracil (m<sup>5</sup>Um) | m<sup>1</sup>A: m<sup>5</sup>Um H:WT |
| 2'-O-methyl uracil (Um) | Um:A W:WC, Um:A<br>W:HT, Um:G H:SC |
| 4-thiouridine (s<sup>4</sup>U) | s<sup>4</sup>U:A W:HT, s<sup>4</sup>U:A<br>W:ST, s<sup>4</sup>U:A S:SC |

<sup>a</sup>Methylated bases are represented as m<sup>x</sup>N, where 'm' represents methyl group, the superscript 'X' represents the position of the methyl group and 'N' represents the nucleobase. m<sup>x</sup><sub>y</sub>N is used to represent more than one methyl substituent, where the subscript 'Y' represents the number of methyl substituents. 2'-O-ribose methylation is represented as Nm.

**TABLE 2.**
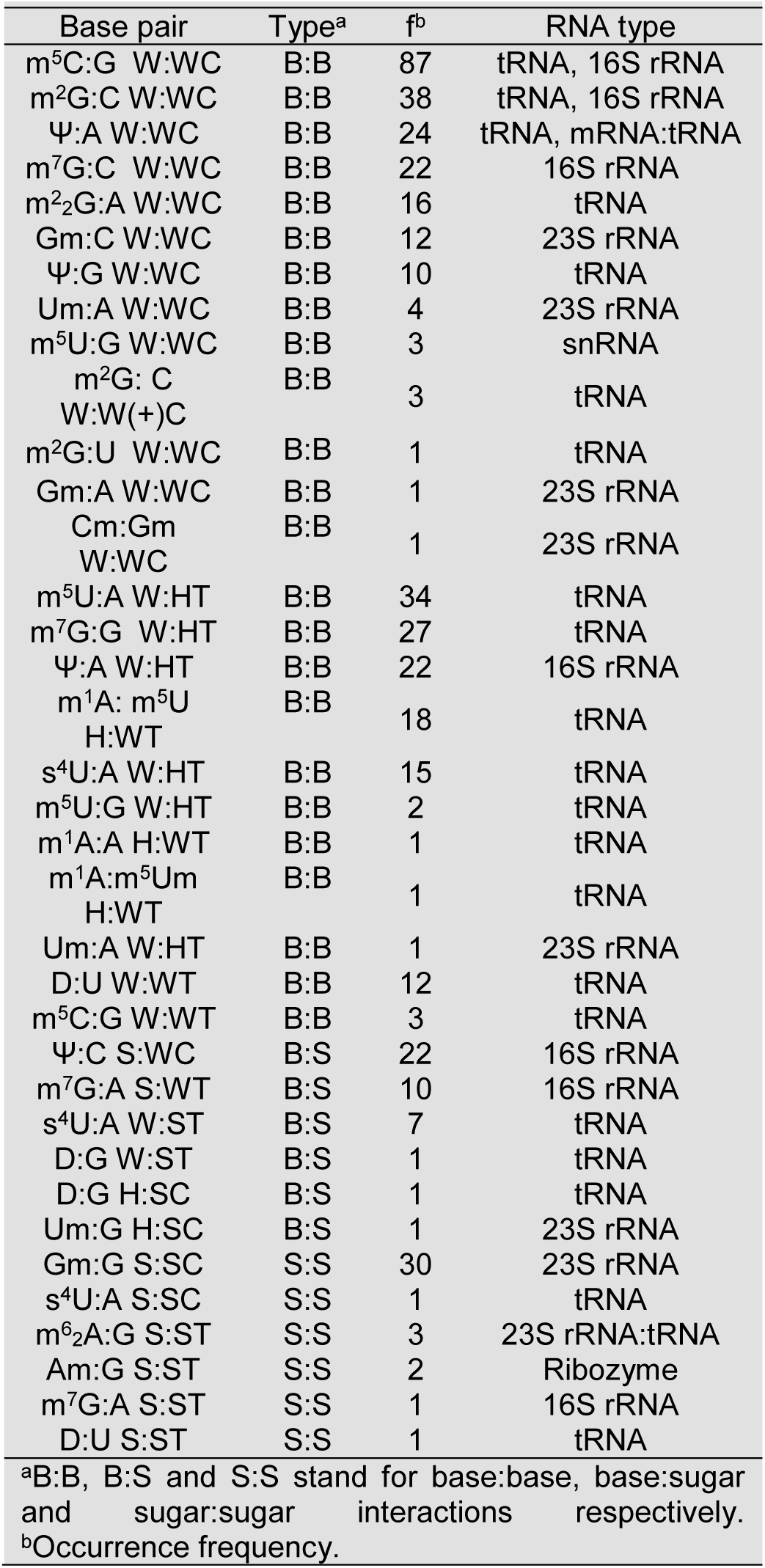
Occurrence frequency and the type of RNA in which the 36 unique modified base pairs were identified in the dataset.

#### Statistical analysis of modified base pairs in tRNA sequence database

As mentioned above, the greatest fraction of modified base pairs are observed in tRNA crystal structures. Owing to the greater number of available tRNA sequences compared to 3D structures, tRNA sequence analysis can provide more detailed information about conservation patterns of modified base pairs. However, the usual method of sequence annotation in the sequence database available at the National Center for Biotechnology Information (Ncbi Resource Coordinators 2015) does not include information on the presence of modified bases in nucleic acid sequences. This excludes the possibility of use of sequence alignment algorithms, such as BLAST (Mládek et al. 2011) etc. for analysis of modified base pairs in the sequence space.

To overcome this difficulty, we have used the tRNA sequence database (Jühling et al. 2009), which is a repository of sequences that provides information on presence of modified bases at different tRNA positions (see Materials and Methods). Within these sequences, base pair combinations present at 10 different positions were searched and graded according to their occurrence in all the sequences (Fig. 4, Supplemental Table S10). These positions were selected due to significant (10 or more) occurrences of modified base pairs at these positions in tRNA crystal structures. Analysis of sequences reveals additional examples of modified base pairs in all the 10 selected positions in the tRNA structure. For example, although the m^2^2G:A combination in W:WC geometry observed most frequently at the 26:44 position in tRNA crystal structures is most frequently observed within the tRNA sequences, our sequence analysis reveals three new modified base pair combinations (m^2^2G:U, m^2^2G:Um and m^2^G:A) at this position. Similarly, at the position 54:58 of TΨC-loop in tRNA, although m^5^U:A W:HTcombinationis the most frequent and covaries with m^5^U:m^1^A, m^5^Um:m^1^A and A:m^1^A pairs in tRNA crystal structures, sequence analysis reveals 5 new modified base pair combinations (m^1^Ψ:A, U:m^1^A, Ψ:m^1^A, Ψ:A and m^5^Um:A) at this position. On similar lines, tRNA sequence analysis identified 6additional modified base pair combinations at other important positions (Fig. 4), which were not observed in tRNA crystal structures.

**FIGURE 4.**
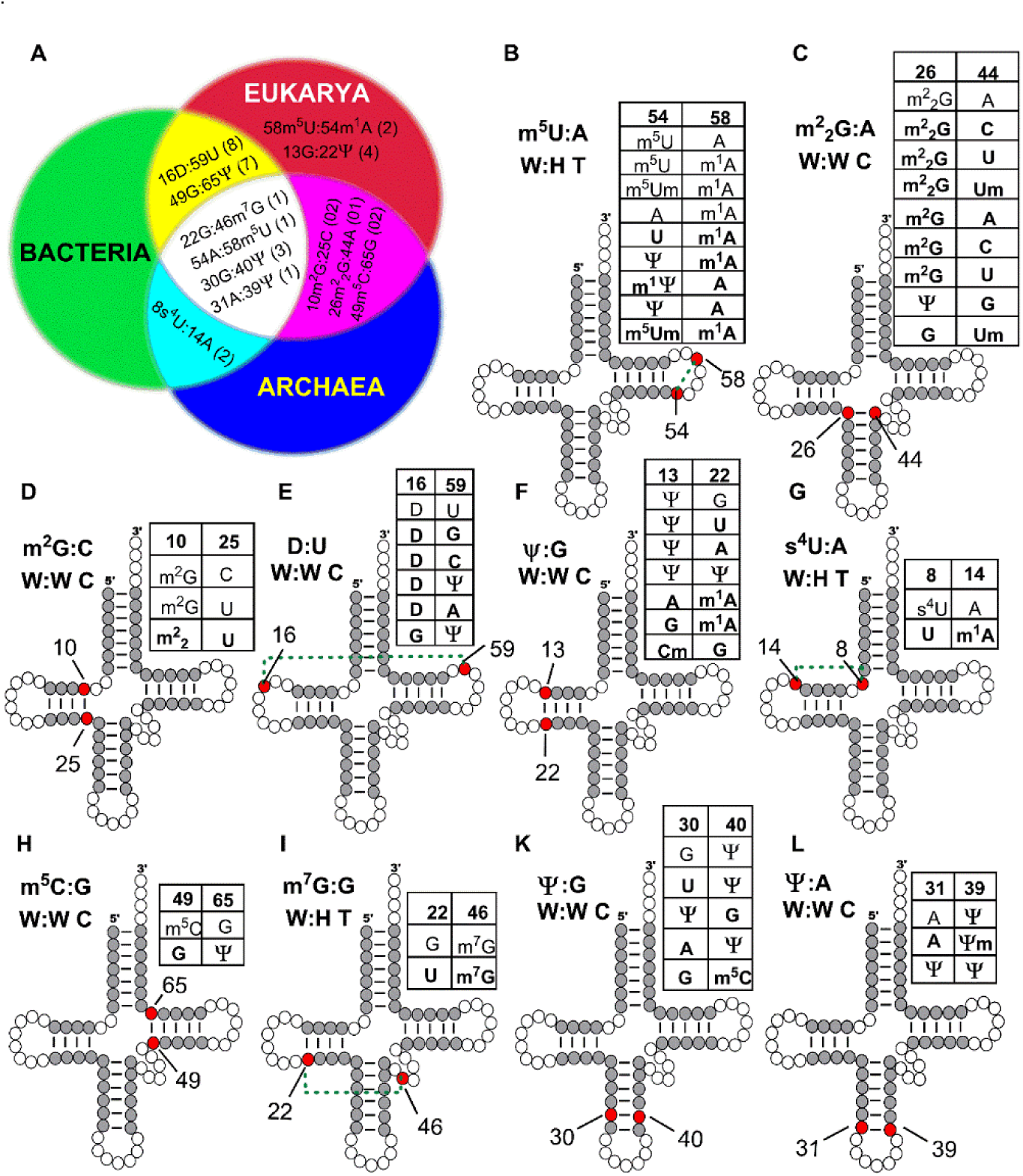
Schematic representation of most commonly observed modified base pairs in tRNA sequences. (A) Distribution of modified base pairs in tRNA sequences divided according to the domains of life. (B-L) Presence of modified base pairs in 10 major base pair positions (represented by red circles) in tRNA structures. The newly identified modified base pair combinations observed from sequence analysis are shown in bold in the corresponding tables.

**FIGURE 5.**
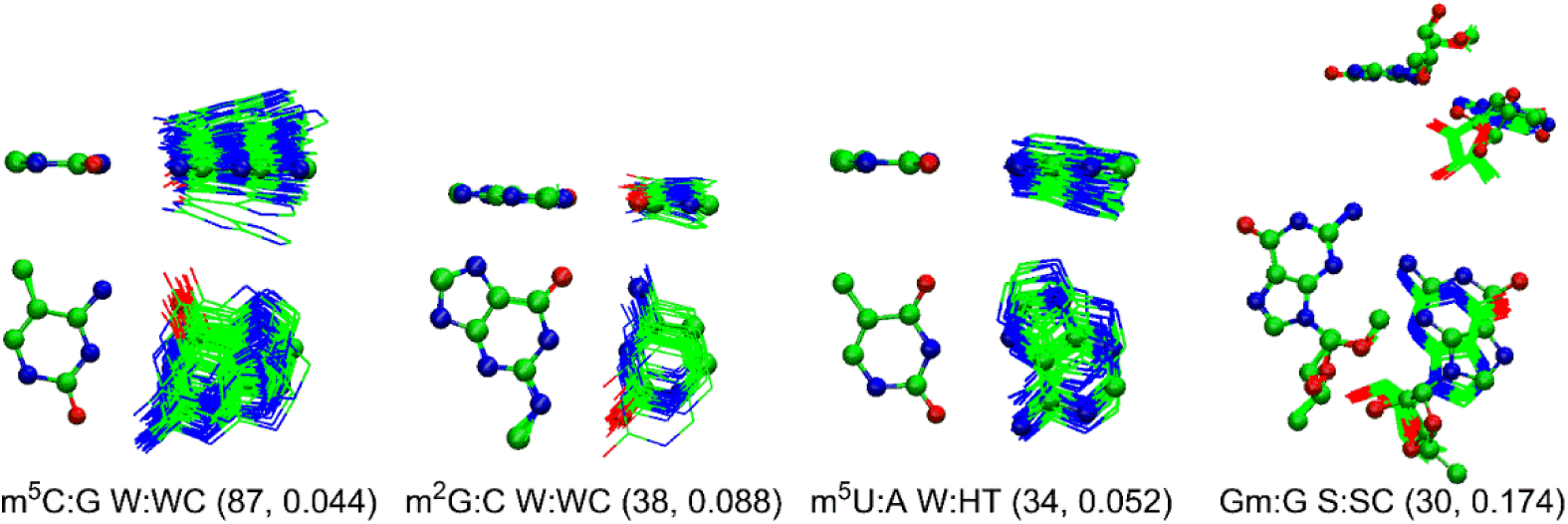
Structural alignment of crystal occurrences of modified base pairs (with occurrence frequency ≥ 30) with their corresponding optimized structures. Occurrence frequency and average RMSD with respect to the optimized structure (in Å) given in the parenthesis.

Our analysis further reveals that some modified base pairs are localized in certain domains of life, and are completely absent in others. For example, although modified base pairs m^2^G:C, m^2^_2_G:A and m^5^C:G at positions10:25, 26:44 and 49:65 respectively are observed in tRNA sequences of archaea and eukarya, and are absent in bacteria (Fig. 4A). Similarly, the m^5^U:m^1^A base pair present at 54:58 position and Ψ:G base pair at 13:22 position was found only in eukaryotic tRNA, and is absent in lower domains (bacteria and archaea). While these examples suggest the absence of some modified base pair combinations in lower organisms, certain modified base pairs are only observed in lower organisms, and have not reached higher domains of life. For example, thiouridine base pair, s^4^U:A at position 8:14 is observed in bacteria and archaea but not in eukarya. Nevertheless, four modified base pairs G:m^7^G, A:m^5^U, G:Ψ and A:Ψ observed at tRNA positions 22:46, 54:58, 30:40 and 31:39 respectively are present in all three domains of life. Overall, our results point towards phylogeny-dependent distribution of modified base pairs in tRNA, which may stem from domain-specific strategies of RNA maturation (Machnicka et al. 2014).

#### Geometric and energetic characterization of modified base pairs

Of the 36 unique modified base pair combinations identified in RNA crystal structures, 23 involve more than one occurrences (Table 2). Geometrical deviations were observed within the multiple occurrences of each base pair. Such deviations arise due to variation in macromolecular context of occurrence of base pairs, and depend on the identity of the base pairs. We used eight different structural parameters, viz. root mean square deviation (rmsd), buckle (κ), propeller twist (π), open angle (σ), stagger (s_x_), shear (s_y_), stretch (s_z_) and E-value to quantify the variation in geometries and hydrogen bonding characteristics of modified base pairs in their crystal contexts (Supplemental Tables S9-S12, Supplemental Section S1 and Supplemental Fig. S2–S5). Analysis of average and standard deviation in these parameters reveals that most of the base pairs involving B-B interactions exhibit relatively smaller deviation among crystal occurrences. However, significant variation is observed in base pairs involving B-S and S-S interactions, which can be mainly attributed to the flexibility of ribose sugar and association glycosidic torsional freedom.

Geometry optimization of a crystal occurrence of each modified base pair using quantum chemical methods allowed us to locate the minimum energy structures of isolated base pairs. These optimized isolated base pair structures represent the ideal base geometries that would be obtained in the absence of macromolecular crystal structure effects, and are useful to quantify the role of interbase hydrogen bonding in determining the structure of the pair. Thus the comparison of geometries of the base pairs observed in its isolated form, those observed in RNA structural context, can provide useful insights into the interplay of the forces within the crystal environment. Our results reiterates that the variation in geometrical parameters and E-values between the crystal and the optimized geometry of each base pair depend on the geometry of the base pair, the type of interaction (B:B vs. B:S vs. S:S) and the identity of the interacting bases. For example, whereas the high rmsd_av_ (0.8 Å) of the optimized structure of Um:A W:WC pair compared to its crystal occurrences is because of significant relaxation of buckle and propeller parameters on optimization, the high rmsdav (1.2 Å) of the optimized structure of m^2^_2_G:A W:WC from its crystal occurrences can be explained in terms of optimization of hydrogen bond distances (and consequent large deviation in E-values) on optimization. Detailed comparison of structural parameters in crystal geometries and energy minimized (optimized) geometries of the base pairs are provided in Supplemental Section S1.

Comparison of the optimized geometries of modified base pairs with their unmodified counterparts can reveal important clues regarding the effect of base modification on the geometries of base pairs. In this context, geometrical deviations between each modified base pair and its unmodified counterpart was measured and analyzed (Supplemental Section S1, Tables S9-S12 and Figs. S2-S5). Comparison of geometries and interaction energies of modified and unmodified base pairs is expected to provide the influence of base modification on the structure and strength of hydrogen bonding interactions between the pairing bases (Supplemental Table S13). Based on our analysis, the effects of base modification on base pairing can be divided into two broad categories:

i. *Base pairs where modification induces electronic effects*: These include 17 base pairs that involve significant (>2 kcal/mol) change in interaction energy on base modification. Such base pairs can further be grouped into five subcategories:
  a. *Base pairs involving alteration of charge on modification*: These include 7 base pairs, 4 of which belong to W:HT family (m^7^G:G, m^5^U:m^1^A, m^5^Um:m^1^A and A:m^1^A) and one each to W:WC (m^7^G:C), S:WT (m^7^G:A) and S:ST (m^7^G:A) families. All of these acquire positive charge on modification. The resulting enhanced electrostatic component of interaction energy significantly increases the overall base pairing energy (by up to 15 kcal/mol) on modification.
  b. *Base pairs involving alteration of hydrogen bonding on modification*: These include two base pairs, *viz*. Ψ:C S:WC and Gm:G S:SC. The former base pair disrupts one of the interbase hydrogen bonding, resulting in decrease in binding energy by 6.3 kcal/mol. However, the later base pair, Gm:G S:SC alters the H-bonding interactions without affecting the stability of the base pair.
  c. *Base pairs involving change in position of electronegative atoms on modification*: These include three base pairs involving Ψ. Since, Ψ differs from U only in terms of direction of glycosidic bond (e.g. a *trans* base pair involving U will be equivalent to a corresponding cis pair involving Ψ), replacement of U with Ψ changes the location of glycosidic nitrogen with respect to the partner base. This results in change in binding energy of the base pair (Supplemental Table S14).
  d. *Base pairs involving replacement of the highly electronegative element (O) with a lesser electronegative element (S) present on the interacting edge*. This category includes the s^4^U:A W:HT base pair, where O4 atom present on the WC edge is replaced by S atom. Since atom S4 is not involved in interbase hydrogen bonding in the modified or the unmodified base pair, the interaction energy of s^4^U:A W:HT is similar (0.4 kcal/mol) to unmodified pair.
  e. *Base pairs involving change in aromaticity of the nucleobase ring on modification*. These include four dihydrouridine containing base pairs, *viz*. D:U W:WT, D:G W:ST, D:G H:SC and D:U S:ST, where the change in interaction energy range between upto 2.8 kcal/mol. Since the loss of aromaticity increases the pucker of the pyrimidine (D) ring, dihydrouridine-containing base pairs adopt different geometries compared to their unmodified counterparts.
ii. *Base pairs where modification may result in alteration of the surrounding steric environment*: This category include 19 base pairs that involve negligible (<2 kcal/mol) change in interaction energy on base modification, indicating that modification does not significantly change the electronic structure of the base pairs. Alternatively, it possible that base modification in such base pairs is important for providing appropriate steric alterations of the local environment within the RNA macromolecular structures. Such alterations may include blocking the hydrogen bonding capability of the base pair with other surrounding nucleosides, or change the conformational space available to other ligands/proteins present at the interface. Depending on the site of modification, such base pairs can further be grouped into two classes:
  a. *Base pairs involving change in steric environment on the minor groove side*. These include 11 base pairs, which involve modification of the amino group of guanine (m^2^G and m^2^_2_G) or the 2’-OH (Am, Cm, Gm and Um). Such modifications may alter the accessibility of the minor groove of the base pair, resulting in potential disruption of associated RNA motifs.
  b. *Base pairs involving change in steric environment on the major groove side*. These include 8 base pairs that involve m^5^C, m^5^U, m^6^_2_A or s^4^U bases, where modification occurs at the major groove side of the base pair. Through introduction of the hydrophobic (methyl) groups on nucleobases, such modifications affect the conformational space available for other molecules such as proteins, ligands, other RNA and antibiotics (Demirci et al. 2014) to interact with RNA.

### Functional roles of modified base pairs

#### Investigating structure – function correlations for some frequently occurring modified base pairs

Although our comprehensive analysis of the RNA crystal structure database, tRNA sequence database and quantum chemical calculations provided useful insights into the occurrence frequencies of modified base pairs within different RNA classes and associated their geometric and energetic features, analysis of macromolecular structural context of occurrence of modified base pairs, as well as their associated functional roles is expected to provide an understanding of “why” base modifications occur in the context of RNA (Supplemental Fig. S7 to S10). Based on several clues from experimental structures available in literature, and adequately supported by our own structural analysis we provide structural and energetic explanations on why nature may have invoked the modification of bases in functional RNA.

i. *Presence of methylated base pairs at the hinge regions of tRNA facilitates molecular flexibility*: Our study reveals that the majority of modified base pairs occur in tRNA, where methylation is the most common modification present in such base pairs. The substantial occurrence of methylated base pairs in tRNA raises question on whether such base pairs are associated with certain specific structural and functional roles in tRNA. It is well known that tRNA is designed to be a flexible molecule for facilitating its dynamic interactions with the ribosome during various stages of translation (Frank et al. 2005). Specifically, the helical stems of tRNA are placed in unique stereo chemical positions during its interaction with ribosome, which are required for the peptide transfer and other tRNA transitions during protein synthesis (Voorhees and Ramakrishnan 2013). Specifically, two tRNA structural regions have previously been proposed to act as hinges for providing flexibility during tRNA transitions (Frank et al. 2005), (Sanbonmatsu 2006). These include the interface of the D-stem/anticodon stem, which further include the base pairing positions 10:25 and 26:44, and the TΨC-stem/acceptor-stem which includes the 49:65 base pairing position (Supplemental Fig. S11). Our analysis reveals the presence of methylated base pairs at both these hinge regions of tRNA. Specifically, the 10:25 position involves three modified base pairs (m^2^G:C, m^2^G:U and m^2^_2_G:U) with substantially high overall occurrence frequency in tRNA sequences, whereas the methylated base pairs present at the 26:44 and 49:65 positions involve six (m^2^_2_G:A, m^2^_2_G:Um, m^2^_2_G:U, m^2^G:A, m^2^G:C, m^2^G:U and m^2^G:Um) and one (m^5^C:G), with significant (16 and 30 respectively) occurrence frequencies in tRNA sequences. Further, analysis of crystal structures examplify the occurrence of a single, but unique (m^2^G:C W:WC at 10:25, m^2^_2_G:A W:WC at 26:44 and m^5^C:G W:WC at 49:65) modified base pair at each of these positions, albeit with significant occurrence frequency (19 at 10:25, 16 at 26:44 and 25 at 49:65). Analysis of base pairing geometries present in these three positions of tRNA reveals that presence of methylated base pairs may help in preventing unnecessary hydrogen bonding interactions involving these base pairs with surrounding bases during tRNA transitions, which in turn provides the required flexibility to the tRNA structure. For example, the presence of m^2^_2_G at 26 position within the 26:44 pair blocks the N2 donor from forming unnecessary hydrogen bonds at this junction (Supplemental Fig. S11). This base pair possesses relatively high average buckle (κ_av_ = 23.4°) and propeller (π_av_ = −22.3°) in crystal occurrences, and possesses significant deviations in buckle within the crystal geometries, compared to the ideal optimized geometries, indicating the inherent geometrical flexibility within is pair. This, coupled with the fact that the interaction energy of the modified base pair is higher than the unmodified counterpart by ~1.6 kcal/mol, indicates that methylation at the 26:44 pair may possibly contribute to the flexibility of the tRNA structure, without compromising the base pair stability. This point towards the potential role of the methyl substituents of the nucleobases in providing flexibility to the tRNA structures, and facilitating its dynamics, adds to the list of previously identified roles of methylated bases, including enhancing stacking interactions, altering conformation equilibrium of ribose sugar (Davis 1998)and altering interbase interactions etc. in context of RNA structure and dynamics (Helm 2006).
ii. *Putative role of sugar methylation in tRNA accommodation on the ribosome platform*: The process of ‘accommodation’ of tRNA on the ribosome is a key conformational change for tRNA selection during translation (Whitford et al. 2010). However, the entropy of free 3’-CCA end of aminoacyl-tRNA opposes accommodation (Fig. 6A), which provides a time delay necessary for the transition of tRNA from A/T to A/A site on the ribosome (Whitford et al. 2010). During this time delay, the momentary, but significant interaction occurs between the 3’–CCA end of tRNA and the A-loop of 23S rRNA (Sanbonmatsu et al. 2005). Specifically, the crystal structure of tRNA:rRNA complex of *H. marismortui* (Nissen et al. 2000) reveals that the C75 base of 3’-CCA end of tRNA is accommodated at the A-loop of helix 92of rRNA (Fig. 6B), where it interacts with Gm2588 in a W:WC base pairing orientation (Fig. 6C). This Gm2588 further interacts with the minor groove of the C2542:G2617 pair in helix 90 of rRNA to form the G-minor base triplet (Nissen et al. 2000; Hansen et al. 2002). Thus, the W:WC interaction of the C75 base of 3’-CCA end of tRNA with the Gm2588 results in the formation of C75 (tRNA) : Gm2588 (H92) : G2617 (H91) : C2542 (H91) quartet at the 3-way junction of H89, H91 and H93 in 23rRNA, which stabilizes the free 3’–CCA end of tRNA during translation process. This 3’–CCA end is further stabilized through stacking interactions involving the A-loop nucleobases of rRNA. Specifically, the quartet C75:Gm2588:G2617:C2542 is flanked on both sides with base triples formed by remaining 3’–CCA bases C74 and C76 with the nucleobases of A-loop and H91. Surprisingly, although this triplet-quartet-triplet association is retained in the analogous crystal structure of E. coli (PDB: 4V9D) in the similar sequence context, the Gm at 2588 is replaced by unmodified G. Our analysis of the E. coli crystal structures reveal that the interaction of 2’–OH group of G2588 with the G2617 reduces the planarity and decreases the hydrogen bond strength within the C75:Gm2588:G2617:C2542 quartet (PDB: 3CME, Fig. 6C). On the other hand, although the presence of Gm at 2588 position in *H. marismortui* alters the H-bonding interactions involving its 2’–OH group with G2617, without affecting the base pair stability, significant optimization of almost all the base pair parameters is observed on sugar methylation (Supplemental TablesS11-S12). Further, the C2’–endo conformation of Gm2588 helps it to undergo rapid *syn-anti* conformational switching on interaction of tRNA with the ribosome, which allows it to interact with the C75 of tRNA with little energetic penalty (Blanchard and Puglisi 2001). Thus it appears that the ribose methylation of G2588 helps in maintaining the interaction of tRNA with large ribosomal subunit and helps in smooth transition of tRNA from A/T phase to A/A phase. Thus, the presence of a modified guanine nucleotide (ribose methylated, Gm2588) in evolutionarily advanced archaea *(H. marismortui)* appears to provide it with a structural advantage over bacteria *(E. coli)* for optimizing the tRNA-rRNA interactions during protein synthesis. This underscores the potential role of modified bases, and corresponding base pairs in facilitating the RNA-RNA interactions in nature.
iii. *Putative role of modified base pairs in T-loop motif of tRNA*: The T-loop of tRNA (commonly known as TΨC loop) represents the prototypic T-loop structural motif (Chan et al. 2013) formed by five consecutive nucleotides (Supplemental Fig. 8A). The TΨC loop is an example of a lone pair triloop motif, since it includes three unpaired residues and a single intra loop base pair at the position 54:58 with non-canonical U:A W:H T geometry, which acts as the loop closing base pair. The backbone environment at the 54:58 pair is so flexible that despite significant differences in C1’-C1’ distances (~9 and ~12 Å respectively), it can accommodate both purine–pyrimidine as well as purine–purine residues with various degrees of modifications. From our tRNA sequence analysis, we observed 9 modified W:HT base pairing combinations (m^5^U:A, m^5^U:m^1^A, m^5^Um:A, m^5^Um:m^1^A, m^5^U:G, m^1^Ψ:A, Ψ:m^1^A, Ψ:A, and A:m^1^A) at the 54:58 position in tRNA., of which 5 (m^5^U:A, m^5^U:m^1^A, m^5^Um:m^1^A, m^5^U:G and A:m^1^A) were also available in tRNA crystal structures. Out quantum chemical analysis suggest that all five modified base pairs observed at 54:58 position in tRNA crystal structure increase the interaction energy (up to 7.4 kcal/mol) compared to their corresponding unmodified counterparts (Supplemental Fig. 8A). This indicates that the presence of modified base pair within the T-loop may provide additional stabilization to the motif. Further, this T-loop is also known to participate in tertiary interactions with the D-loop of tRNA, thus forming base pairs at 18:55 and 19:56 positions that maintain a continuous stack on top of 54:58 base pair and form a mini duplex (Supplemental Fig. 8A). Since base modifications are known to enhance stacking interactions (Agris 1996), it is possible that the modified pairs at 54:58 positions, may play an important role in stabilizing the T-loop, and its associated tertiary interactions with the D-loop, which may in turn help in maintaining the functional conformation of tRNA.
iv. *Possible role of methylated base pairs in the antibiotics binding regions of the bacterial ribosome*: Previous experimental studies on the interaction of aminoglycosidic antibiotics with bacterial ribosome revealed that methyl modification of rRNA bases is known resist the entry and binding of antibiotic compounds in the binding pocket (Demirci et al. 2014). Such modified bases act either by modulating their surrounding hydrophobic environment and consequently affecting the conformational space of antibiotic binding region, or by blocking the entry of these antibiotics into the binding pocket (Demirci et al. 2014). For example, streptomycin and paromomycin are known to induce errors in the mRNA decoding process by affecting the local dynamics of rRNA nucelobases that interact with the codon-anticodon complex and are present in the decoding region at the A-site of 30S subunit of ribosome (Fig. 7). Previous crystal structure studies suggest that streptomycin interacts with the phosphate backbone (Carter et al. 2000) of helices 18, 27 and 44 of 16S rRNA and amino acid residues of S12 protein of the ribosome. Incidentally, N7-methylation at G527 of the m^7^G527:C522 (G:C W:WC) pair of helix 18 in 16S rRNA has previously been shown to cause streptomycin resistance (Demirci et al. 2014). However, the structural role of N7-methylation in inducing streptomycin resistance is not well understood. Our analysis indicates that N7-methylation of the G:C W:WC imparts positive charge to G, which enhances the intrinsic stability of the base pair by ~9 kcal/mol, while maintaining geometry similar to that of the canonical G:C base pair. This indicates that although N7-methylation may change the hydrophobic environment in the antibiotic binding pocket, without destabilizing the G527:C522 base pair. Specifically, m^7^G527 is present only at a distance of ~4.0 Å from the antibiotic binding pocket, where it imparts noticeable structural differences to its surroundings by influencing the position and orientation of hydrophobic side chain of the amino acid residues K46, K47, P48, N49 and K91 of the S12 protein (Fig. 7F), which may in turn affect the backbone conformation and size of the binding pocket. In case of paromomycin, it is known from literature that 5-methylation of C1407 in helix 44 of 16S rRNA causes low resistance towards antibiotic binding in the A-site of 16S rRNA (Demirci et al. 2014). This might be attributed towards steric hindrance induced by methyl substituent on C1407. The crystal structure of 16S rRNA suggests that C1407 forms a W:WC pair with G1494 of helix 44, which is present near the mRNA decoding region. Our analysis indicates C5-methyl modification does not affect the intrinsic stability of the C1407:G1494 base pair, where it maintains a geometry similar to that of canonical C:G W:WC base pair. Since paromomycin interacts with the major groove of base pair C1407:G1494 W:WC only when C1407 is non-methylated (Vicens and Westhof 2001), and provided that methylation does not change the stability of the C1407:G1494 pair, it indicates that the role of methylation lies in providing a steric hindrance for antibiotic binding, without affecting the electronic structure within the binding pocket (Fig. 7E). This illustrates the role of steric effect of modified bases in determining the RNA-ligand interactions.
v. *Potential involvement of modified base pairs in higher order structures, and their putative functional roles*. Our analysis revealed that several modified base pairs are involved in formation of higher order structures such as base triplets and quartets. Specifically, we observed eleven distinct triplets and two quartet structures spanning nine modified bases within the dataset (Supplemental Table S17). Fig. 8 shows the geometrical arrangement of some representative higher order structures that have potential functional roles. Some such recurrent motifs involving protonated base pairs are discussed below:
  a. *Ψ:A:A motif*: The Ψ4:A36:A1493 (PDB: 4JV5) is an mRNA:tRNA:rRNA interaction motif in *T. thermophiles*, and is a part of the codon-anticodon complex at the ribosome decoding region. Similar to the unmodified UAA motif, Ψ:A:A is an A-minor interaction motif (Nissen et al. 2001) composed of Ψ:A W:WC (a first mRNA codon -tRNA anticodon) and A36:A1493 S:SC (tRNA-rRNA) base pairs (Fig. 8). Although the unmodified U4 is the first alphabet in (UAA or UAG or UGA) mRNA stop codons, its replacement with Ψ4 changes the stop codons to sense codons, where they code for serine/threonine (ΨAA or ΨAG) and tyrosine/phenylalanine (ΨGA) amino acids, which provides a new way to expand the genetic code (Fernandez et al. 2013).
  b. *m^5^U:G:A motif*: This is A-minor interaction is observed at the 5’-splice site of group-I self-splicing intron in *Azoarcus* sp. (Adams et al. 2004). This wobble receptor motif is composed of a wobble pair between m^5^U1 at the end of the exon and G10 within the internal guide sequence. This G10:m^5^U1 W:WC pair is recognized from its minor groove by the conserved A58 base present within the A-rich loop of the intron, where it forms the G10:A58 S:SC interaction (PDB: 1U6B, Fig. 8).
  c. *D:G:C motif*: This motif represents the loop-loop tertiary interaction involving the D loop and variable loop at the elbow region of tRNA, which is composed of G15:C48 W:WT and D20:G15 W:ST base pairs (PDB 1SER, Fig. 8). In this motif, exocyclic amino group of G15 base pair forms bifurcated base pair, where it simultaneously forms H-bonds with both D20 (O2) and C48 (N3).
  d. *Gm:G:C triplet and C:Gm:G:C quartet motifs*: In the *H. marismortui* 23S rRNA (PDB 3CME), Gm2588 present in the A-loop of helix 92 folds back on helix 90 and interacts with the nucleotides C2542 and G2617, and forms a Gm2588:G2617:C2542 triplet (Hansen et al. 2002). This triplet consists of C2542:G2617 W:WC and Gm2588:G2617 S:SC pairs, which represent a unique G-minor motif where the Gm2588 interactions from the minor grove of the canonical G:C W:WC base pair. During the process of tRNA accommodation at the A-site of the ribosome, this motif interacts with the C75 of 3’-CCA end of tRNA (Nissen et al. 2000), and forms a quartet (Fig. 8). Crystal structure analysis suggests that the quartet motif is further stabilized by two flanking base triplets involving C74 and A76 of 3’-CCA end of A-site tRNA.
  e. *G:Um:A motif*: This motif is a part of the sarcin-ricin domain (Correll et al. 2003) situated in helix 95 in domain VI of large ribosomal subunit of *E.coli*, and includes the most prevalent GpU dinucleotide platform interaction (Lu et al. 2010). This motif is composed of both B-B (Um2656:A2665 W:HT) and B-S (Um2656:G2655 H:SC) interactions (PDB 3DW5, Fig. 8). Although the GpU platform nucleotides showed single strong H-bond interaction between U2656 (O4) and G2655 (N2), additional stabilization is provided by base phosphate interactions. Methylation at 2’-OH of U2565 blocks the minor grove interactions without affecting other interactions within the motif.
  f. *s4U:A:A motif*: This motif is observed in *E.coli* tRNA^Cys^ and formed when s^4^U8 from acceptor/D stem junction in tRNA interacts with the D loop bases A14 and A21, forming B-B (s^4^U8:A14 W:HT) and B-S (s^4^U8:A21 S:WC) interactions (Fig. 8). The U8:A14 interaction is particularly important for the formation of the core of the tRNA structure by bringing the D loop in closer proximity for base pairing with residues in the T loop (Lauhon et al. 2004). Apparently, this motif is conveniently opportunistic, since the third base A21 is sometimes replaced by neighboring base A46 with S:SC geometry instead of W:SC (PDB 1B23, Supplemental Fig. S7).

**FIGURE 6.**
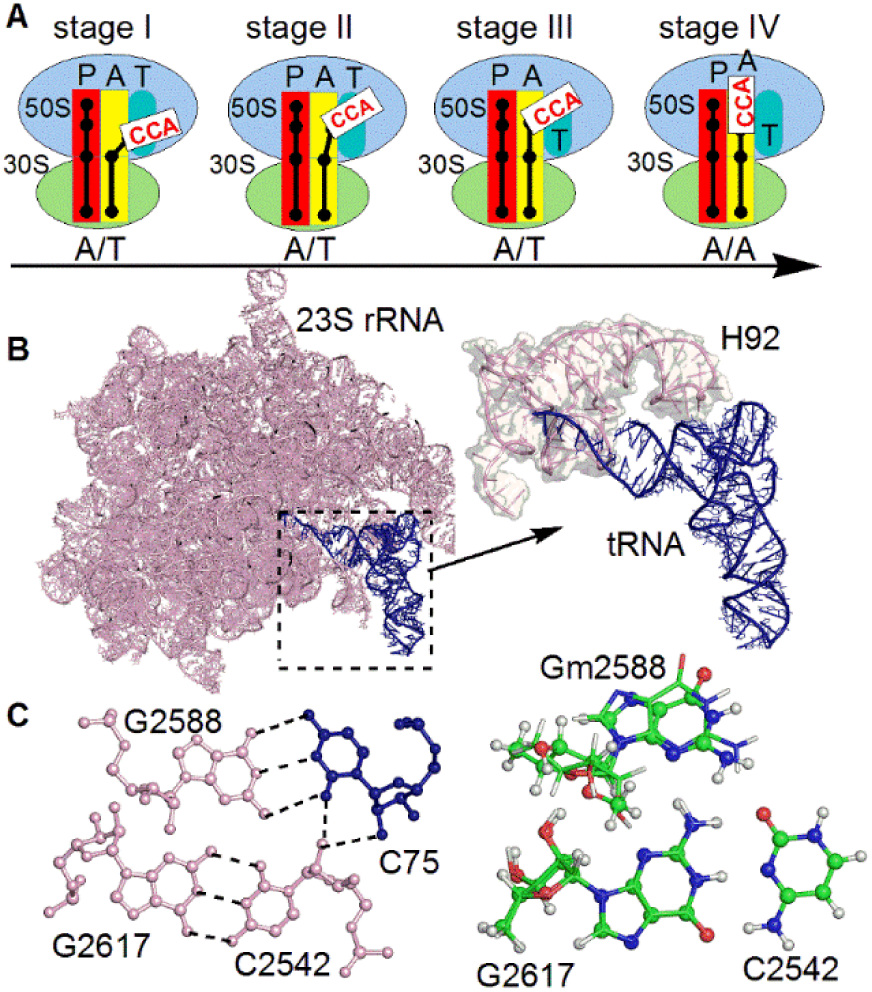
(A) Flexible 3’–CCA end (white box) of tRNA during various stages of tRNA accommodation at the A-site (yellow box) of the 70S ribosome. The neighboring P-site of rRNA is shown as red box. (B) Interaction of 3’–CCA containing amino acceptor arm of tRNA blue) of tRNA (blue) with the A-loop (H92) of 23S rRNA (pink). (C) Structure of base quartet formed from the interaction of the preformed G-minor base triplet (C2542:G2617:Gm2588) present at H92 of rRNA and the C75 of the 3′–CCA of tRNA. Alignment of the preformed rRNA triplet containing the 2’–methylated G2588 present in the crystal structure of tRNA:rRNA complex of *H. marismortui* (PDB: 3cme), with the corresponding triplet containing the unmodified G2588 present in one of the crystal structure of tRNA:rRNA complex of *E.coli* (PDB: 4v9d).

**FIGURE 7.**
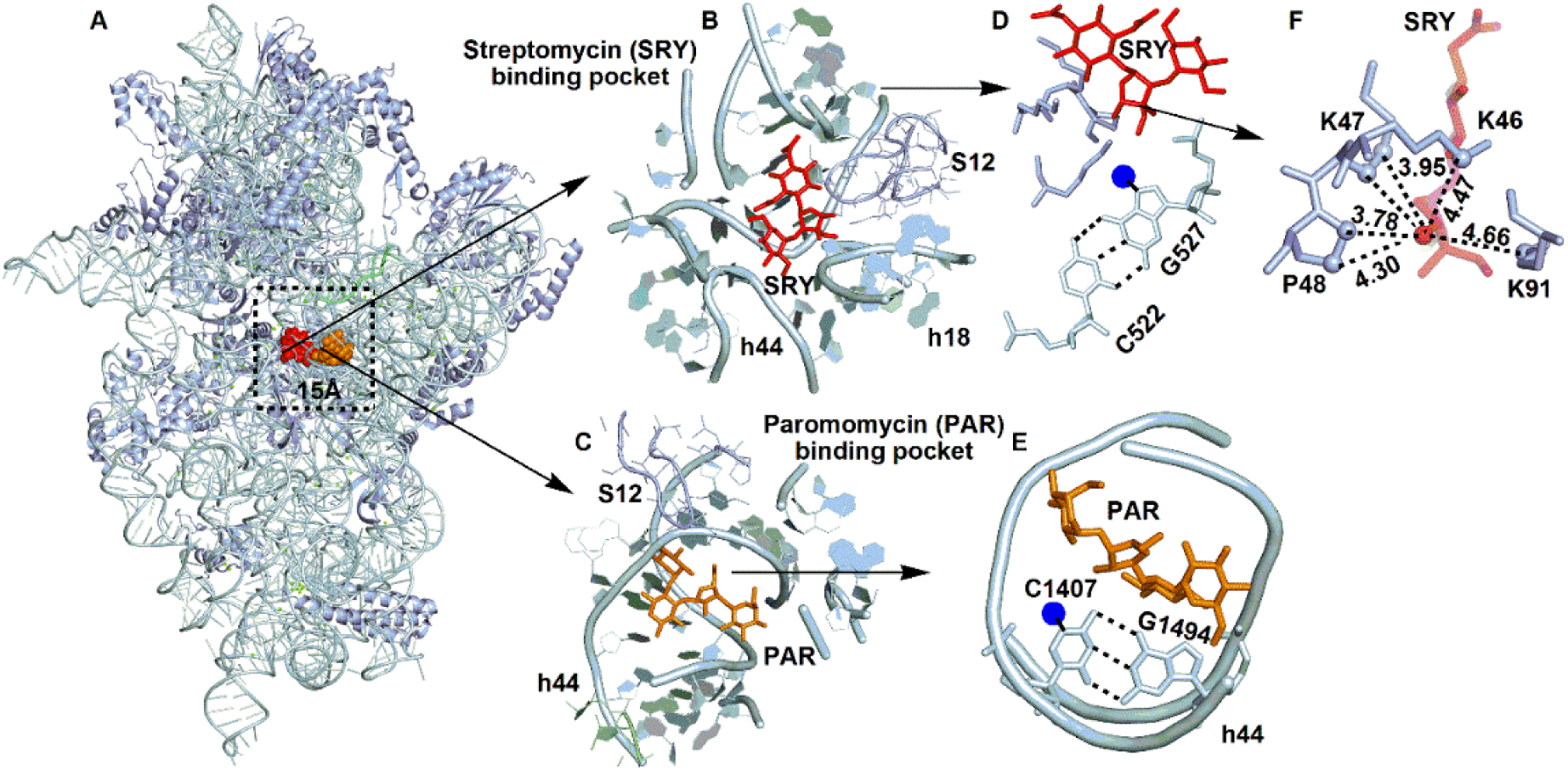
Presence of modified base pairs at the binding site of antibiotics streptomycin and paromomycin. (A) Structure of 16S rRNA bound to streptomycin (red) and paromomycin (orange) (B, C) Antibiotic binding pocket with surrounding proteins (S12). (D, E) Interaction of base pairs C522:527 and C1407:G1494 present in the binding pocket with the antibiotic streptomycin and paromomycin respectively. (F) Hydrophobic cloud created by surrounding amino acid residues around the methyl group attached to ring II of streptomycin (red). Methyl modification of G527 or C1407 at the nucleobase sites represented by blue circles result in resistance to antibiotic binding.

**FIGURE 8.**
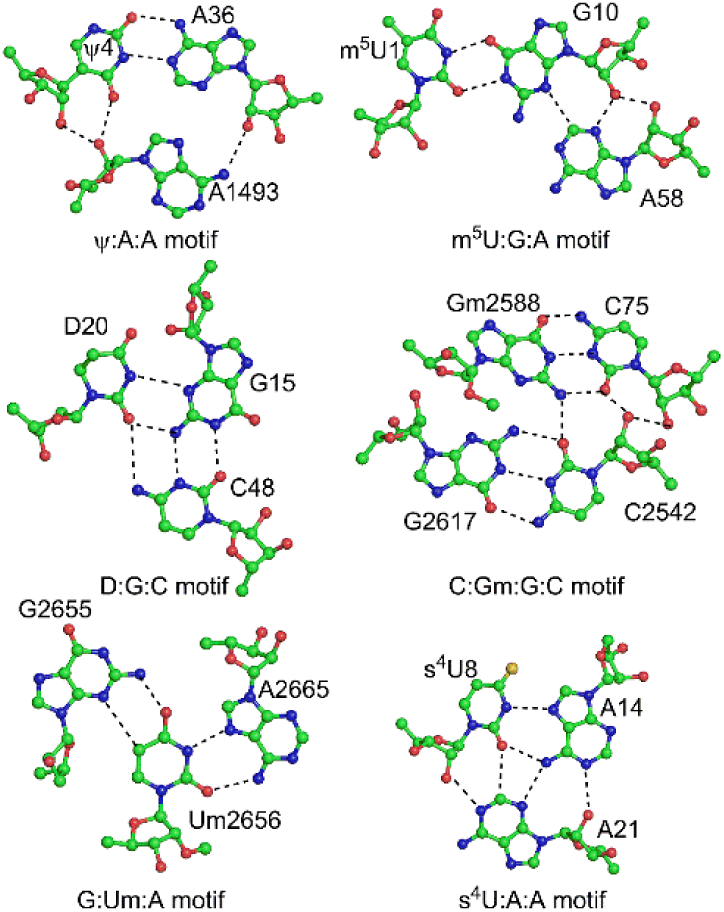
Modified base pairs involved in higher order interaction motifs.

## CONCLUSIONS

We carried out detailed statistical, geometrical, energetic and contextual analysis of 36 naturally occurring post-transcriptionally modified base pairs present in RNA macromolecules. Such base pairs span diverse structures and include base-base, base-sugar and sugar-sugar interactions. Our results reveal that although overall greater proportion of modified base pairs occurs in tRNA, base pairs containing sugar modifications are conspicuously more abundant in rRNA. Further analysis of available tRNA sequences reveals 28additional examples of modified base pairs at 10 selected positions in the tRNA structure that are not observed in RNA crystal structures. This adds to the available list of modified base pairs, and underscores the importance of sequence analysis in understanding of conservation patterns of RNA motifs.

In general, methylated base pairs are found to be more abundant compared to base pairs containing other modifications in RNA, which can be correlated to the variety of functional roles that methylated bases play in functional RNA, including altering base conformation, affecting base stacking, etc. Further, base pairs containing uracil and guanine modifications are more abundant compared to those containing modifications of cytosine or adenine, which can be explained on the basis of the occurrence of substantial variety in types of uracil modifications and guanine methylations in RNA. Detailed analysis of local RNA topology around the location of modified base pairs in RNA reveals that such base pairs are present in almost all major RNA motifs, which include stems (helices, 49%), loops (14%) and tertiary interactions (37%), and point to the diverse structural roles that modified base pairs may play in RNA.

We used advanced quantum chemical methods (MP2/aug-cc-pVDZ//B3LYP/6-31G (d,p) to analyze the optimal geometries, strength of inter base interaction and effect of base modification on the geometries and energies of RNA base pairs. Using examples of base pairs containing uracil and guanine modifications, we illustrated the effect of type and position of chemical modification on the geometries and stabilities of base pairs. On the basis of change in strength of interaction on base modification, the effects of base pair modification were further classified into steric and electronic perturbations on the unmodified base pairing geometry. Analysis of surrounding macromolecular environment as well as local RNA structural topology around the modified base pairs revealed certain important structural and functional contexts, including those involving unique modified base pairs in tRNA, as well as sugar modified base pairs in rRNA, which suggest that some of them may be playing important roles in maintaining the structure, dynamics and functions of RNA molecules. Overall, our highlight the prevalence of modified base pairs in RNA, and indicate that greater level of understanding role of these interacting motifs in many biological processes involving RNA is yet to be achieved.

## MATERIALS AND METHODS

### Dataset of RNA crystal structures

To identify base pairs containing modified bases, the occurrence of modified bases was first search in RNA crystal structures. For this, the PDB*sum* (Laskowski 2009) database, which summarizes information on X-ray crystal structures deposited in the protein databank (PDB) was used. Specifically, using the ‘Het Groups’ option of PDB*sum*, a unique 3-letter code corresponding to each of the 15 modified residues (Table 1) was used to retrieve the relevant list of PDB entries submitted till 18 July 2016. The retrieved crystal structures were further filtered according to their resolution, and in synchrony with previous crystal structure study (ref.), structures with resolution better than 3.5 Å were selected for further analysis. The dataset is intentionally kept redundant with respect to sequence, since the previous study has shown that possible modified base pair types and base conformations may differ within crystal structures of same RNA (Chawla et al. 2015). BPFind software (Das et al. 2006) was used to analyze the occurrence, location and type of modified base pairs with at least two hydrogen bonds in the selected RNA crystal structures.

### tRNA sequence analysis

We analyzed all the 474 tRNA cytoplasmic sequences belonging to 73 organisms (i.e. prokaryotes (19), archaea (9), eukaryotes (41) and viruses (4)) from the tRNAdb database (Jühling et al. 2009). For each sequence, we recorded which bases are present at positions where modified base pairs occur in the analyzed crystal structures of tRNA. Thus, at each of the position, the relative occurrence frequency of the modified base pair was recorded, and ranked within all available combinations. Once the tRNA sequences that contained modified base pairs at specific positions were identified, the sequences that contained the modified pair were further classified according to the type of corresponding aminoacyl tRNA.

### Quantum mechanical energy minimization and interaction energies

Among 36 distinct base pair combinations studied, 24 combinations contained base pairs with only base-base interactions, 6 combinations contained base-nucleoside interactions and 6 base pairs contained sugar-sugar interactions. For geometry optimization (energy minimization) of the base pairs that do not involve interaction ribose sugar with the pairing base, the C1’ atoms of both the participating nucleosides were replaced with hydrogen atoms. For the base pairs involving base-nucleoside interactions, depending on whether one or both the sugars are involved in base pairing, the respective ribose sugars were retained during energy minimization. In these cases, the 5’–OH group of the interacting ribose sugar was replaced by hydrogen atom, whereas the 3’–OH group was retained during calculations.

Geometry optimization of the base pairs was carried out at the B3LYP/6-31G (d,p) (Lee et al. 1988; Becke 1993) level using Gaussian 09 (Frisch et al. 2009), which was selected in synchrony with previous studies on RNA base pairs (Šponer et al. 2004; Šponer et al. 2005d; Sharma et al. 2008; Sharma et al. 2010b). The strength of hydrogen bonds between two bases of the modified base pair was calculated in terms of binding energy or interaction energy, which is defined as the extra stabilization acquired by two bases when the form the base pair. Thus, the interaction energy (ΔE^AB^) of a base pair AB composed of two bases A and B is defined as

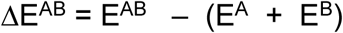

where E^AB^ is energy of the base pair and E^A^ and E^B^ are the energies of bases A and B respectively. Interaction energies were calculated at the RIMP2/aug-cc-pVDZ level (Weigend and Häser 1997; Ahlrichs et al. 1998), which was selected in analogy with previous studies. The interaction energies were corrected for basis set superposition error (Boys and Bernardi 1970) using the Turbomole v6.2 (Karlsruhe and GmbH 2011) suite of quantum chemical programs.

### Comparison of macromolecular crystal and optimized geometries of base pairs

#### RMSD

To understand the difference in base pair geometry in optimized form and in crystal occurrences, the root mean square deviations (RMSD) of crystal constrained geometries of each modified base pair was calculated from their corresponding optimized geometries. In addition, to analyze the variation in base pair geometry within crystal occurrences, the RMSD of each crystal occurrence of the base pair was calculated with respect to the average structure among all crystal occurrences. These calculations were done using VMD v1.9 software (Humphrey et al. 1996).

#### Base-pair parameters

Change in the geometries of base pairs upon optimization, and variation in structures of crystal occurrences of base pairs were quantitatively evaluated by comparing the base pair parameters (buckle, propeller, open angle, shear, stretch and stagger) of the crystal occurrences with the optimized geometry of each base pair, as well as among the different crystal occurrences of the base pair. These calculations were done using upgraded version of NUPARM software (Bansal et al. 1995), which uses the edge-specific system for calculation of base pair parameters, which is specific to RNA base pairs.

#### E-Values of hydrogen bonds

To evaluate the relative goodness of hydrogen bonds within base pairs in their crystal occurrences as well as in optimized geometries, we have calculated a parameter called E-value, which is defined as:

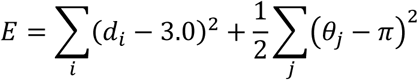

Here d is the heavy atom distance for each hydrogen bond between two bases under consideration and θ is a pseudo angle subtended by precursor atoms of both the bases (Das et al. 2006). This parameter was used, since the RNA crystal structures from which base pairs were extracted, did not contain hydrogen atoms. The E-value parameter assess the quality of hydrogen bonds in the absence of hydrogen atom coordinates, and is useful in analyzing hydrogen bonds within the crystal occurrences of base pairs.

## REFERENCES

Adams PL, Stahley MR, Kosek AB, Wang J, Strobel SA. 2004. Crystal structure of a self-splicing group I intron with both exons. Nature 430: 45–50.

Agris PF. 1996. The Importance of Being Modified: Roles of Modified Nucleosides and Mg2+ in RNA Structure and Function. In Progress in Nucleic Acid Research and Molecular Biology, Volume 53 (ed. EC Waldo, M Klvle), pp. 79–129. Academic Press.

Ahlrichs R, Elliott SD, Huniar U. 1998. Quantum chemistry: Large molecules — small computers. Berichte der Bunsengesellschaft für physikalische Chemie 102: 795–804.

Bansal M, Bhattacharyya D, Ravi B. 1995. NUPARM and NUCGEN: software for analysis and generation of sequence dependent nucleic acid structures. CABIOS 11: 281–287.

Becke AD. 1993. Density-functional thermochemistry. III. The role of exact exchange. J. Chem. Phys. 98: 5648–5652.

Blanchard SC, Puglisi JD. 2001. Solution structure of the A loop of 23S ribosomal RNA. Proc. Natl. Acad. Sci. USA 98: 3720–3725.

Boys SF, Bernardi F. 1970. The calculation of small molecular interactions by the differences of separate total energies. Some procedures with reduced errors. Mol. Phys. 19: 553–566.

Cantara WA, Crain PF, Rozenski J, McCloskey JA, Harris KA, Zhang X, Vendeix FAP, Fabris D, Agris PF. 2011. The RNA modification database, RNAMDB: 2011 update. Nucleic Acids Res. 39: D195–D201.

Carter AP, Clemons WM, Brodersen DE, Morgan-Warren RJ, Wimberly BT, Ramakrishnan V. 2000. Functional insights from the structure of the 30S ribosomal subunit and its interactions with antibiotics. Nature 407: 340–348.

Chan CW, Chetnani B, Mondragón A. 2013. Structure and function of the T-loop structural motif in noncoding RNAs. Wiley Interdisc. Rev.: RNA 4: 507–522.

Charette M, Gray MW. 2000. Pseudouridine in RNA: What, Where, How, and Why. IUBMB Life 49: 341–351.

Chawla M, Oliva R, Bujnicki JM, Cavallo L. 2015. An atlas of RNA base pairs involving modified nucleobases with optimal geometries and accurate energies. Nucleic Acids Res. 43: 9573.

Chawla M, Sharma P, Halder S, Bhattacharyya D, Mitra A. 2011. Protonation of Base Pairs in RNA: Context Analysis and Quantum Chemical Investigations of Their Geometries and Stabilities. J. Phys. Chem. B 115: 1469–1484.

Correll CC, Beneken J, Plantinga MJ, Lubbers M, Chan YL. 2003. The common and the distinctive features of the bulged-G motif based on a 1.04 Å resolution RNA structure. Nucleic Acids Res. 31: 6806–6818.

Dalluge JJ, Hashizume T, Sopchik AE, McCloskey JA, Davis DR. 1996. Conformational flexibility in RNA: the role of dihydrouridine. Nucleic Acids Res. 24: 1073–1079.

Das J, Mukherjee S, Mitra A, Bhattacharyya D. 2006. Non-Canonical Base Pairs and Higher Order Structures in Nucleic Acids: Crystal Structure Database Analysis. J. Biomol. Struct. Dyn. 24: 149–161.

Davis DR. 1998. Biophysical and Conformational Properties of Modified Nucleosides in RNA (Nuclear Magnetic Resonance Studies). In Modification and Editing of RNA. American Society of Microbiology.

Decatur WA, Fournier MJ. 2002. rRNA modifications and ribosome function. Trends Biochem. Sci. 27: 344–351.

Demirci H, Murphy FV, Murphy EL, Connetti JL, Dahlberg AE, Jogl G, Gregory ST. 2014. Structural Analysis of Base Substitutions in Thermus thermophilus 16S rRNA Conferring Streptomycin Resistance. Antimicrob. Agents Chemother. 58: 4308–4317.

Denmon AP, Wang J, Nikonowicz EP. 2011. Conformation Effects of Base Modification on the Anticodon Stem–Loop of Bacillus subtilis tRNATyr. J. Mol. Biol. 412: 285–303.

Dunin-Horkawicz S, Czerwoniec A, Gajda MJ, Feder M, Grosjean H, Bujnicki JM. 2006. MODOMICS: a database of RNA modification pathways. Nucleic Acids Res. 34: D145–D149.

Emmerechts G, Maes L, Herdewijn P, Anné J, Rozenski J. 2008. Characterization of the Posttranscriptional Modifications in Legionella pneumophila Small-Subunit Ribosomal RNA. Chem. Biodiv. 5: 2640–2653.

Fernandez IS, Ng CL, Kelley AC, Wu G, Yu Y-T, Ramakrishnan V. 2013. Unusual base pairing during the decoding of a stop codon by the ribosome. Nature 500: 107–110.

Frank J, Sengupta J, Gao H, Li W, Valle M, Zavialov A, Ehrenberg M. 2005. The role of tRNA as a molecular spring in decoding, accommodation, and peptidyl transfer. FEBS Lett. 579: 959–962.

Frisch MJ, Trucks GW, Schlegel HB, Scuseria GE, Robb MA, Cheeseman JR, Scalmani G, Barone V, Mennucci B, Petersson GA et al. 2009. Gaussian 09. Gaussian, Inc., Wallingford, CT, USA.

Halder A, Bhattacharya S, Datta A, Bhattacharyya D, Mitra A. 2015. The role of N7 protonation of guanine in determining the structure, stability and function of RNA base pairs. Phys. Chem. Chem. Phys. 17: 26249–26263.

Halder A, Halder S, Bhattacharyya D, Mitra A. 2014. Feasibility of occurrence of different types of protonated base pairs in RNA: a quantum chemical study. Phys. Chem. Chem. Phys. 16: 18383–18396.

Hansen MA, Kirpekar F, Ritterbusch W, Vester B. 2002. Posttranscriptional modifications in the A-loop of 23S rRNAs from selected archaea and eubacteria. RNA 8: 202–213.

Helm M. 2006. Post-transcriptional nucleotide modification and alternative folding of RNA. Nucleic Acids Res. 34: 721–733.

Humphrey W, Dalke A, Schulten K. 1996. VMD: visual molecular dynamics. J. Mol. Graph. Model. 14: 33–38.

Jühling F, Mörl M, Hartmann RK, Sprinzl M, Stadler PF, Pütz J. 2009. tRNAdb 2009: compilation of tRNA sequences and tRNA genes. Nucleic Acids Res. 37: D159–D162.

Karlsruhe Uo, GmbH F. 2011. Turbomole 6.2.

Laskowski RA. 2009. PDBsum new things. Nucleic Acids Res. 37: D355–D359.

Lauhon CT, Erwin WM, Ton GN. 2004. Substrate Specificity for 4-Thiouridine Modification in Escherichia coli. J. Biol. Chem. 279: 23022–23029.

Lee C, Yang W, Parr RG. 1988. Development of the Colle-Salvetti correlation-energy formula into a functional of the electron density. Phys. Rev. B 37: 785–789.

Leontis NB, Stombaugh J, Westhof E. 2002. The non-Watson–Crick base pairs and their associated isostericity matrices. Nucleic Acids Res. 30: 3497–3531.

Leontis NB, Westhof E. 2001. Geometric nomenclature and classification of RNA base pairs. RNA 7: 499–512.

Limbach PA, Crain PF, McCloskey JA. 1994. Summary: the modified nucleosides of RNA. Nucleic Acids Res. 22: 2183–2196.

Lu X-J, Olson WK, Bussemaker HJ. 2010. The RNA backbone plays a crucial role in mediating the intrinsic stability of the GpU dinucleotide platform and the GpUpA/GpA miniduplex. Nucleic Acids Res. 38: 4868–4876.

Machnicka MA, Olchowik A, Grosjean H, Bujnicki JM. 2014. Distribution and frequencies of post-transcriptional modifications in tRNAs. RNA Biol. 11: 1619–1629.

Mládek A, Sharma P, Mitra A, Bhattacharyya D, Šponer J, Šponer JE. 2009. Trans Hoogsteen/Sugar Edge Base Pairing in RNA. Structures, Energies, and Stabilities from Quantum Chemical Calculations. J. Phys. Chem. B 113: 1743–1755.

Mládek At, Šponer JE, Kulhánek P, Lu X-J, Olson WK, Šponer Ji. 2011. Understanding the sequence preference of recurrent RNA building blocks using quantum chemistry: the intrastrand RNA dinucleotide platform. J. Chem. Theory Comput. 8: 335–347.

Motorin Y, Helm M. 2010. tRNA Stabilization by Modified Nucleotides. Biochemistry 49: 4934–4944.

Mueller EG, Buck CJ, Palenchar PM, Barnhart LE, Paulson JL. 1998. Identification of a gene involved in the generation of 4-thiouridine in tRNA. Nucleic Acids Res. 26: 2606–2610.

Ncbi Resource Coordinators. 2015. Database resources of the National Center for Biotechnology Information. Nucleic Acids Res. 43: D6–D17.

Nissen P, Hansen J, Ban N, Moore PB, Steitz TA. 2000. The Structural Basis of Ribosome Activity in Peptide Bond Synthesis. Science 289: 920.

Nissen P, Ippolito JA, Ban N, Moore PB, Steitz TA. 2001. RNA tertiary interactions in the large ribosomal subunit: The A-minor motif. Proc. Natl. Acad. Sci. USA 98: 4899–4903.

Oliva R, Cavallo L, Tramontano A. 2006. Accurate energies of hydrogen bonded nucleic acid base pairs and triplets in tRNA tertiary interactions. Nucleic Acids Res. 34: 865–879.

Oliva R, Tramontano A, Cavallo L. 2007. Mg2+ binding and archaeosine modification stabilize the G15–C48 Levitt base pair in tRNAs. RNA 13: 1427–1436.

Sanbonmatsu KY. 2006. Alignment/misalignment hypothesis for tRNA selection by the ribosome. Biochimie 88: 1075–1089.

Sanbonmatsu KY, Joseph S, Tung C-S. 2005. Simulating movement of tRNA into the ribosome during decoding. Proc. Natl. Acad. Sci. USA 102: 15854–15859.

Sharma P, Chawla M, Sharma S, Mitra A. 2010a. On the role of Hoogsteen:Hoogsteen interactions in RNA: Ab initio investigations of structures and energies. RNA 16: 942–957.

Sharma P, Mitra A, Sharma S, Singh H, Bhattacharyya D. 2008. Quantum Chemical Studies of Structures and Binding in Noncanonical RNA Base pairs: The Trans Watson-Crick:Watson-Crick Family. J. Biomol. Struct. Dyn. 25: 709–732.

Sharma P, Šponer JE, Šponer J, Sharma S, Bhattacharyya D, Mitra A. 2010b. On the Role of cis Hoogsteen:Sugar Edge Family of Base Pairs in Platforms and Triplets—Quantum Chemical Insights into rNa Structural Biology. J. Phys. Chem. B 114: 10234–10234.

Šponer J, Jurečka P, Hobza P. 2004. Accurate Interaction Energies of Hydrogen-Bonded Nucleic Acid Base Pairs. J. Am. Chem. Soc. 126: 10142–10151.

Sponer J, Šponer JE, Petrov AI, Leontis NB. 2010. Quantum chemical studies of nucleic acids: can we construct a bridge to the RNA structural biology and bioinformatics communities? J. Phys. Chem. B 114: 15723–15741.

Šponer JE, Leszczynski J, Sychrovský V, Šponer J. 2005a. Sugar edge/sugar edge base pairs in RNA: stabilities and structures from quantum chemical calculations. J. Phys. Chem. B 109: 18680–18689.

Šponer JE, Špačková Na, Kulhánek P, Leszczynski J, Šponer J. 2005b. Non-Watson-Crick base pairing in RNA. quantum chemical analysis of the cis Watson-Crick/sugar edge base pair family. J. Phys. Chem. A 109: 2292–2301.

Šponer JE, Špačková Na, Leszczynski J, Šponer J. 2005c. Principles of RNA Base Pairing: Structures and Energies of the Trans Watson–Crick/Sugar Edge Base Pairs. J. Phys. Chem. B 109: 11399–11410.

Šponer JE, Špačková Na, Leszczynski J, Šponer J. 2005d. Principles of RNA base pairing: structures and energies of the trans Watson-Crick/sugar edge base pairs. J. Phys. Chem. B 109: 11399–11410.

Stombaugh J, Zirbel CL, Westhof E, Leontis NB. 2009. Frequency and isostericity of RNA base pairs. Nucleic Acids Res. 37: 2294–2312.

Vicens Q, Westhof E. 2001. Crystal Structure of Paromomycin Docked into the Eubacterial Ribosomal Decoding A Site. Structure 9: 647–658.

Voorhees RM, Ramakrishnan V. 2013. Structural Basis of the Translational Elongation Cycle. Ann. Rev. Biochem. 82: 203–236.

Walsh CT, Garneau-Tsodikova S, Gatto GJ. 2005. Protein Posttranslational Modifications: The Chemistry of Proteome Diversifications. Angew. Chem. Int. Ed. 44: 7342–7372.

Weigend F, Häser M. 1997. RI-MP2: first derivatives and global consistency. Theor. Chem. Acc. 97: 331–340.

Whitford PC, Geggier P, Altman RB, Blanchard SC, Onuchic JN, Sanbonmatsu KY. 2010. Accommodation of aminoacyl-tRNA into the ribosome involves reversible excursions along multiple pathways. RNA 16: 1196–1204.

Yoshizawa S, Fourmy D, Puglisi JD. 1999. Recognition of the codon-anticodon helix by ribosomal RNA. Science 285: 1722–1725.

Yu F, Tanaka Y, Yamashita K, Suzuki T, Nakamura A, Hirano N, Suzuki T, Yao M, Tanaka I. 2011. Molecular basis of dihydrouridine formation on tRNA. Proc. Natl. Acad. Sci. USA 108: 19593–19598.

